# MTM: a multi-task learning framework to predict individualized tissue gene expression profiles

**DOI:** 10.1101/2022.10.19.512838

**Authors:** Guangyi He, Maiyue Chen, Yingnan Bian, Ence Yang

**Affiliations:** Department of Medical Bioinformatics, School of Basic Medical Sciences, Peking University, Beijing 100191, P. R. China; School of Artificial Intelligence, Peking University, Beijing 100191, P. R. China; Enlight Medical Technologies (Shanghai) Co., Ltd, Shanghai 201318, P. R. China

**Author notes:** These authors contributed equally: Guangyi He, Maiyue Chen.

**Keywords:** multi-task learning, deep learning, artificial neural network, transcriptome, cross-tissue prediction

## Abstract

Predicting tissue expression profiles from peripheral ‘surrogate’ samples, especially blood transcriptome, has become an effective alternative when invasive procedures are not ideal. However, existing approaches ignore tissue-shared intrinsic relevance, inevitably limiting predictive performance. Here, we propose a unified deep learning-based multi-task learning framework, Multi-tissue Transcriptome Mapping (MTM), enabling the prediction of individualized expression profiles from any available tissue of an individual. By jointly leveraging individualized cross-tissue information through multi-task learning, MTM achieves superior sample-level and gene-level performance. With the high prediction accuracy and the ability to preserve individualized biological variations, MTM could facilitate both fundamental and clinical biomedical research.

## Background

Gene expression profiles in diverse tissues act as molecular mediators between genotypes and phenotypes and provide a snapshot of the systemic physiological and pathological status of an individual [1]. Deciphering tissue-derived gene expression not only provides insights into the fundamental molecular mechanisms of biological processes [2-4], but also aids in clinical diagnosis, subtyping and management [5-7]. However, tissue transcriptome-based biological investigations and clinical evaluations highly depend on invasive tissue biopsy procedures, which are risky, harmful, and even unavailable in some cases. Thus, mapping biological information from peripheral ‘surrogate’ samples to tissue transcriptomic profiles has become an emerging solution attracting extensive research interest.

Various methods have been developed to predict tissue-specific expression for specific individuals. Early attempts utilized genotypes with the rationale that individual variance in genetically regulated expression (GReX) components of tissue-specific gene expression could be explained by genotypes to some extent, i.e., expression quantitative trait loci (eQTL). A representative work demonstrating this strategy is the PrediXcan, which takes advantage of eQTL variants to construct gene-based tests to impute GReX [8]. However, these eQTL-based methods are limited to genes with significant tissue-specific eQTL single-nucleotide polymorphisms and neglect non-GReX factors in expression. As blood is involved in the circulation and bidirectional exchange of substances with organs, it is considered to reflect the in-time functional status of tissues underlying gene expression, including components attributable to feedback of traits, and to environmental and other factors [9, 10]. Recent studies have supported the advantages of predicting tissue expression profiles based on blood expression [11-14]. Nevertheless, the methods based on genotype and transcriptome usually have two common limitations: first, they are typically based on linear models or traditional machine learning methods, inevitably limiting their capability to capture complex nonlinear expression relationships in biological organisms [15, 16]; second, they build prediction models independently for a single gene in each target tissue, failing to utilize the intrinsic cross-tissue biological inherence underlying transcriptomes of multiple tissues from the same individual, which could be rescued by borrowing information from similar tissues [17-19].

Deep learning has shown its powerful capability for handling high-dimensional, complex, and nonlinear biological data [20-27]. Multi-task learning, which imposes regularity or shared representation by learning tasks simultaneously, is ideal for utilizing the intrinsic shared features of different domains (e.g., different tissues from the same individual) [28-31]. Thus, we developed MTM, a deep-learning-based multi-task learning framework to predict individualized tissue gene expression profiles using any available tissues from a person. MTM integrates mappings among multiple tissues into a single model, enabling the exploitation of the underlying complex nonlinear patterns of tissue gene expression levels within the same individual. With its outstanding predictive power and the ability to capture individualized physiological and pathological variations, MTM demonstrates its potential to accelerate transcriptome-based research and clinical applications.

## Results

### Framework overview

To achieve individualized gene expression prediction, we proposed a deep learning framework, Multi-tissue Transcriptome Mapping (MTM), a unified multi-task learning approach capable of predicting tissue-specific gene expression profiles using any available tissues from an individual (Fig. 1). Gene expression levels of 19,291 protein-coding genes across 49 tissues from 948 individuals in the Genotype-Tissue Expression (GTEx) datasets [32] were used to develop our model. The expression levels of input tissue samples across different tissues were processed by a unified encoder network equipped with the tissue conditioning module (TCM), to encourage personalized representations in a shared latent space. Then, the latent codes were mapped to multiple tissue-specific expression spaces through a prediction task conditioning generator to achieve personalized prediction of tissue expression profiles. Two auxiliary networks, a mapping network and a discriminator network, were employed to smooth the latent space and to refine the prediction results, respectively (see details in the Methods). With the multi-task learning architecture, MTM is not limited to predicting the expressions of a single target tissue based on a single input tissue, but rather supports arbitrary predefined inputs and predicting targets in a unified model, which introduces more flexibility along with potentially improved performance.

**Fig. 1.**
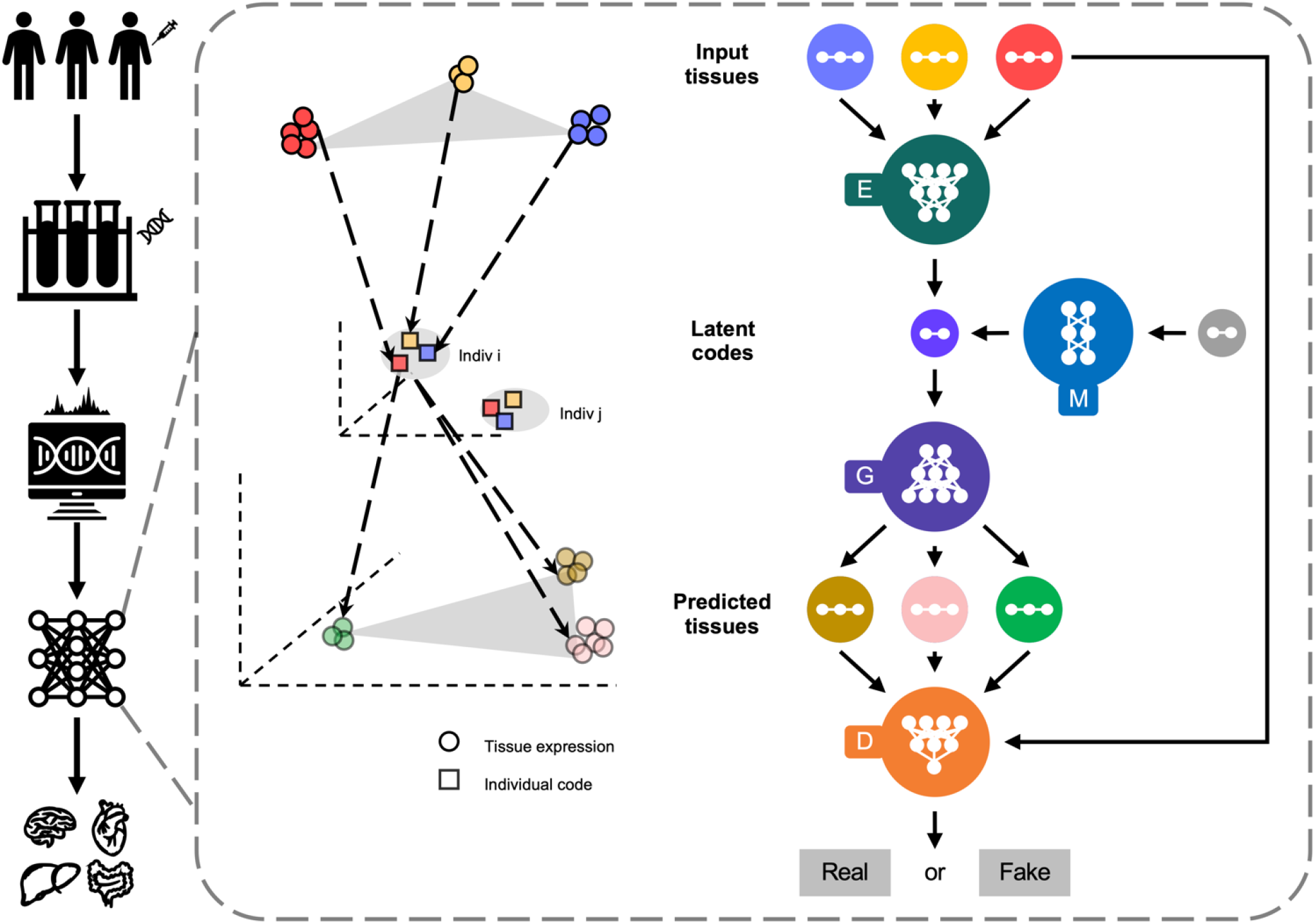
Overview of MTM. MTM is based on a multi-task learning architecture and consists of four artificial neural networks, a unified encoder (*E*), a unified generator (*G*), a unified discriminator (*D*), and a mapping network (*M*). The gene expression profiles of different input tissues are embedded into a shared latent space by *E* to form individualized codes, which are then mapped to multiple tissue-specific expression spaces by *G* to realize personalized prediction of the expression profiles of different target tissues.

### MTM outperforms the baseline methods

The prediction quality of MTM at the sample level and the gene level was evaluated on the task of predicting individualized gene expression profiles of the other 48 tissues within the development set (GTEx) based on the expression profile of whole blood. Two representative single-tissue approaches, B-GEX (method S1) and TEEBoT (method S2), were used as baselines in the evaluation for comparison. At the sample level, the sample-wise prediction accuracy was measured by Pearson’s correlation coefficient, which was 0.21±0.07 across all genes among the 48 tissues (ranging from 0.04 to 0.34). Compared with 0.00±0.02 for S1 and 0.08±0.09 for S2, MTM showed significantly higher levels of accuracy (Wilcoxon rank sum test, *P* < 0.05) than S1 or S2 did in all 48 tissues. On the gene-scale, MTM (0.23±0.07, ranging from 0.04 to 0.34) also demonstrated significantly higher gene-wise accuracy (Wilcoxon rank sum test, *P* < 0.05) than S1 (0.00±0.03) and S2 (0.07±0.11) did in 48 and 47 tissues out of the 48 tissues, respectively (Fig. 2a and Supplementary Table 1).

**Fig. 2.**
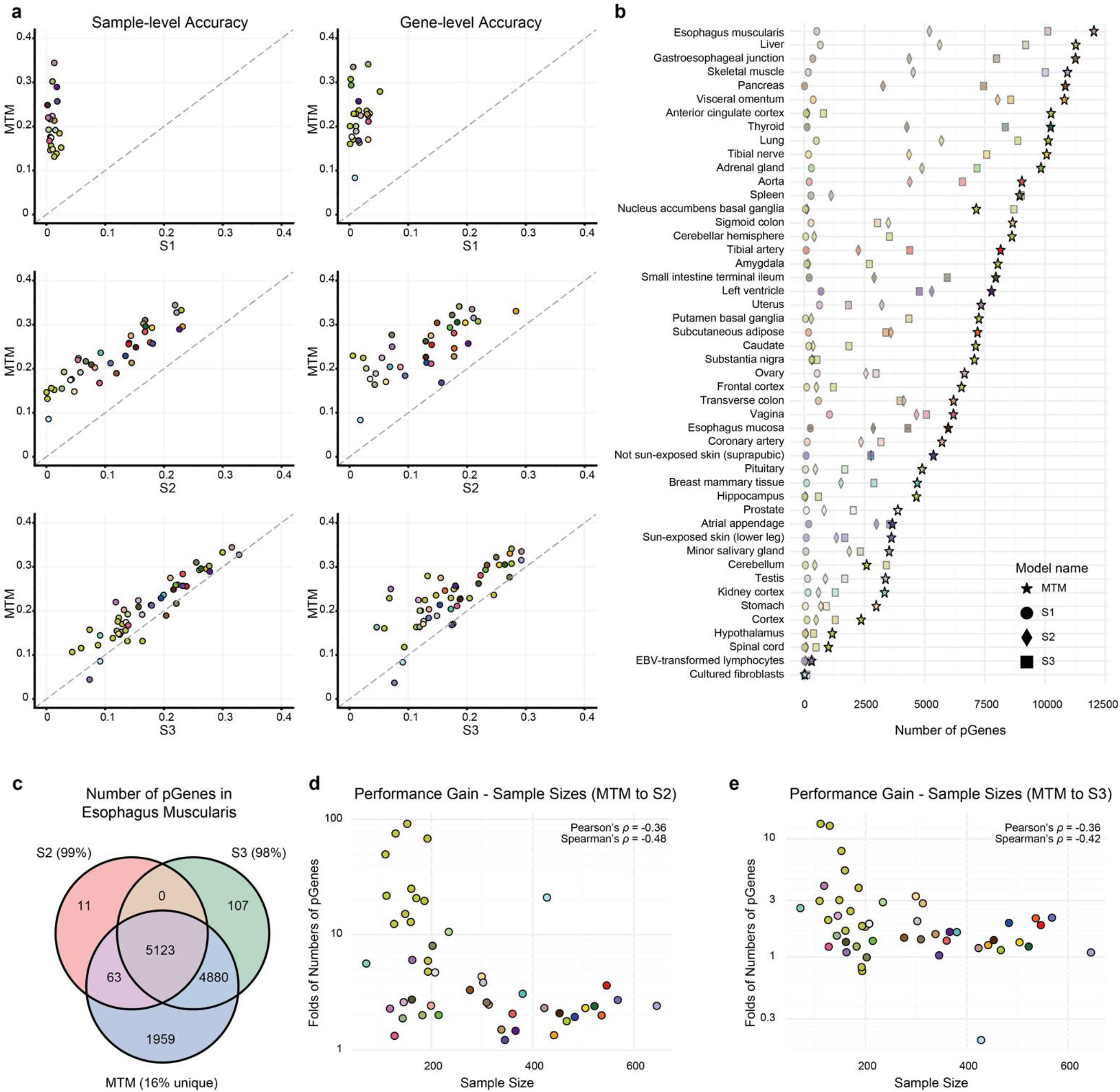
Comparison of the performance for predicting tissue-specific gene expression. **a** The sample-wise and gene-wise accuracy (Pearson’s correlation coefficient) of MTM against single-tissue approaches, including linear (S1 and S2) and nonlinear (S3) models. **b** The number of pGenes (Pearson’s *ρ* > 0.3) of different models in different target tissues. **c** Venn diagram showing the overlap of the pGenes from S2, S3 and MTM in the oesophagus muscularis. Approximately 99% and 98% of pGenes from S2 and S3 were included in the MTM-derived pGenes, respectively. **d, e** The relationship between the fold changes in the number of pGenes and the sample sizes of different tissues for MTM to S2 (**d**) and MTM to S3 (**e**). Each scatter point colour represents a specific tissue as defined by the GTEx Consortium [32].

In terms of predictable genes (pGenes; defined as genes with Pearson’s *ρ* > 0.3 between predicted and observed expression), MTM on average identified 6,597 pGenes across the 48 tissues (ranging from 21 to 12,025), which was substantially higher than the 218 (ranging from 0 to 1,048) of S1 and the 2,301 (ranging from 1 to 8,031) of S2 (Fig. 2b and Supplementary Table 2). For the S1- and S2-derived pGenes, a median of 94.2% (7.7% – 100.0%) and 94.5% (0.0% – 99.8%) were identified by MTM, respectively. In contrast, a median of 2.2% (0.0% – 15.9%) and 32.6% (0.0% – 73.1%) of MTM-derived pGenes were captured by S1 and S2, respectively. When comparing the fold changes in the number of pGenes of MTM to S2 (ranging from 1.22- to 91.37-fold, mean = 11.39-fold, median = 2.91-fold), we found that the performance gain was negatively correlated with sample size (Spearman’s *ρ* = −0.48, *P* = 5.26×10^−4^, Fig. 2d), implying that the missing information caused by small sample sizes could be partially recovered from other tissues with the application of MTM.

### Both the nonlinearity and multi-task framework contribute to performance

To investigate the independent contribution of the nonlinear and multi-task components, we performed ablative experiments and assessed their influence on the overall performance. We constructed an intermediate model (S3) for comparison, which consisted of a set of nonlinear neural networks for each tissue based on simple multilayer perceptrons. For the nonlinearity component, we compared the performance of the linear models (S2) with the constructed nonlinear models (S3) for each tissue. As expected, the accuracy of S3 outperformed S2 both at the sample (0.17±0.07) and gene levels (0.17±0.07). The sample-wise accuracies and gene-wise accuracies of S3 were significantly (*P* < 0.05) higher than those of S2 in 46 tissues and 43 tissues, respectively. Compared with S2, the average number of pGenes resulting from S3 (on average 4,057 pGenes, ranging from 106 to 10,110, Supplementary Table 2) was substantially higher, with fold changes ranging from 0.57- to 106.00-fold (mean = 7.11-fold, median = 1.93-fold).

For the multi-task learning component, MTM was compared with the constructed nonlinear single-tissue models (S3). At the sample level, the accuracy of MTM outperformed S3 in 41 tissues, with the performance in 24 tissues showing statistical significance (*P* < 0.05). In addition, at the gene level, MTM significantly (*P* < 0.05) outperformed S3 in 42 tissues (Fig. 2a and Supplementary Table 1). Furthermore, the average number of pGenes resulting from MTM (6,597) was substantially higher than that of S3 (4,057) (Fig. 2b and Supplementary Table 2), with fold changes ranging from 0.20- to 13.38-fold (mean = 2.46-fold, median = 1.65-fold). Approximately 92.4% of the S3-derived pGenes were still captured by MTM (Fig. 2c and Supplementary Fig. 1). The above results suggested that apart from advantages introduced by nonlinear neural networks, jointly leveraging the individualized cross-tissue information provides extra prediction improvements perpendicularly.

### Intermediate features and outputs of MTM provided insights into the performance basis

To determine how MTM integrated different tissues from the same individual as an entirety, we next explored the characteristics of the individualized representations learned by MTM. Intraindividual latent codes derived from the same individual showed significantly (*P* < 2.2×10^−16^) higher similarities than interindividual latent codes did (Fig. 3b and 3c), indicating that individualized properties of different tissues from the same individual were captured well by MTM. Then, we explored the characteristics of the decoding paths from the same individual representation to the tissue expression profiles. We found that the pairwise similarities between the decoding paths of the data flow (i.e., activations of different layers) were highly consistent (Spearman’s *P* < 0.05) with those between tissue expression profiles (Fig. 3d), suggesting that the biological similarities between tissues were reflected in the decoding process. Together, the integration of multiple tissues into a single model by our multi-task learning architecture resulted in individualized representations and decoding rules with biological patterns.

**Fig. 3.**
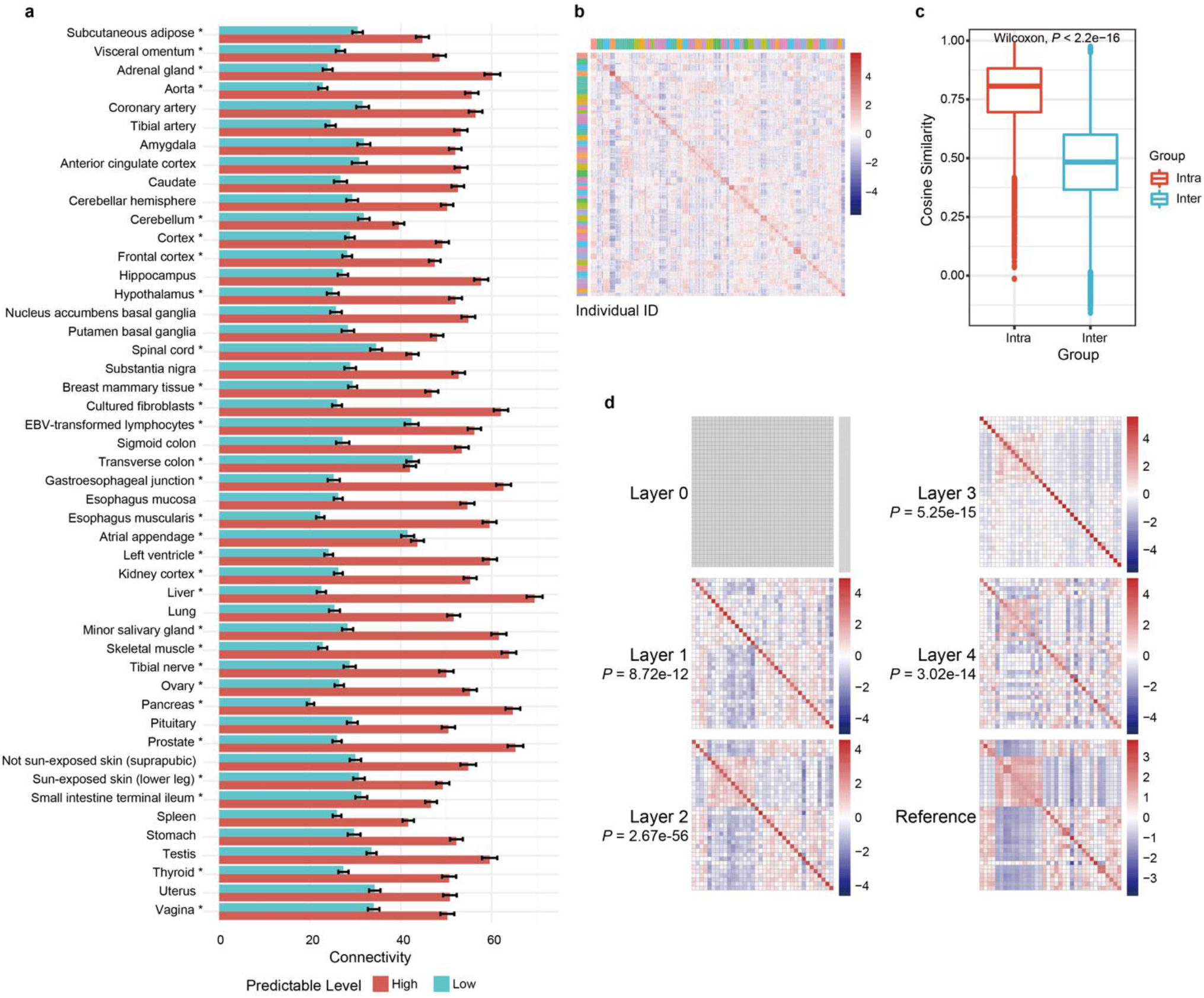
Characteristics of predicted and intermediate data derived from MTM. **a** Comparison of the connectivity levels (average degrees) between highly predictable (top 25 percentiles of predictable genes) and unpredictable genes (bottom 25 percentiles of predictable genes) in different tissues. **b** Heatmap displaying the pairwise similarities of codes of different tissues from 50 random individuals. **c** Comparison of similarities of intraindividual and interindividual latent codes. **d** Pairwise similarities between different intermediate data flow towards different target tissues, which started from the same latent code of the blood expression from one specific individual. Each grid is coloured to indicate a similarity level between intermediate data of the corresponding two tissues. ‘Reference’ refers to pairwise similarities between the expression profiles of different target tissues.

Given that MTM is able to identify more pGenes, distinctive features of the pGenes may also aid in the interpretation of the overall prediction strength of MTM from a different perspective. We found that tissues with similar expression profiles tended to share similar pGenes (Spearman’s *ρ* = 0.56, *P* = 1.40×10^−92^), which was consistent with previous discoveries. The pGenes tended to have significantly (Wilcoxon rank sum test, *P* < 0.05) higher expression levels than the unpredictable genes in all 48 tissues (Supplementary Fig. 3). The pGenes showed significantly (Wilcoxon rank sum test, *P* < 0.05) higher conservation than the unpredictable genes in 9 tissues but significantly lower conservation in 35 out of the 48 tissues (Supplementary Fig. 4), suggesting that there was no obvious preference for conservation in pGenes. Notably, there were significantly higher levels of connectivity (average degrees) in the protein interaction network (PIN) for pGenes than for unpredictable genes in a large portion of tissues (higher in 47 tissues, with 28 tissues showing statistical significance, while significantly lower in one tissue, Wilcoxon rank sum test) (Fig. 3a). By comparison, the S2-derived pGenes showed significantly lower levels of connectivity in 11 tissues, while exhibiting significantly higher levels in 26 tissues. The difference between MTM and S2 suggested that MTM might be more capable of capturing the complex interactions in the PIN.

### MTM could preserve personalized biological variations

As MTM resulted in highly personalized representations of individuals, we assessed the extent to which the predicted tissue expression profiles could reveal trait-associated variations. Correlation analysis of gene expression levels of pGenes was conducted against individuals’ traits (including age, gender and BMI) in the predicted data (from blood expression) and compared with the actual data (i.e., observed data). The age-specific gene expression associations were well preserved (Pearson’s *ρ* > 0.3, *P* < 0.05) in the predicted expression data in 46 out of the 48 tissues, with a *ρ*_*median*_ of 0.81. Similarly, the test statistics of gender (*ρ*_*median*_ = 0.76) and BMI (*ρ*_*median*_ = 0.56) of the predicted data showed significant positive (Pearson’s *ρ* > 0.3, *P* < 0.05) correlations with those of the actual data in all 43 gender-independent tissues and 39 out of the 48 tissues, respectively (Fig. 4a and Supplementary Fig. 5–7). The above results suggested that the individualized tissue expression profiles predicted by MTM preserved the trait-related expression changes well.

**Fig. 4.**
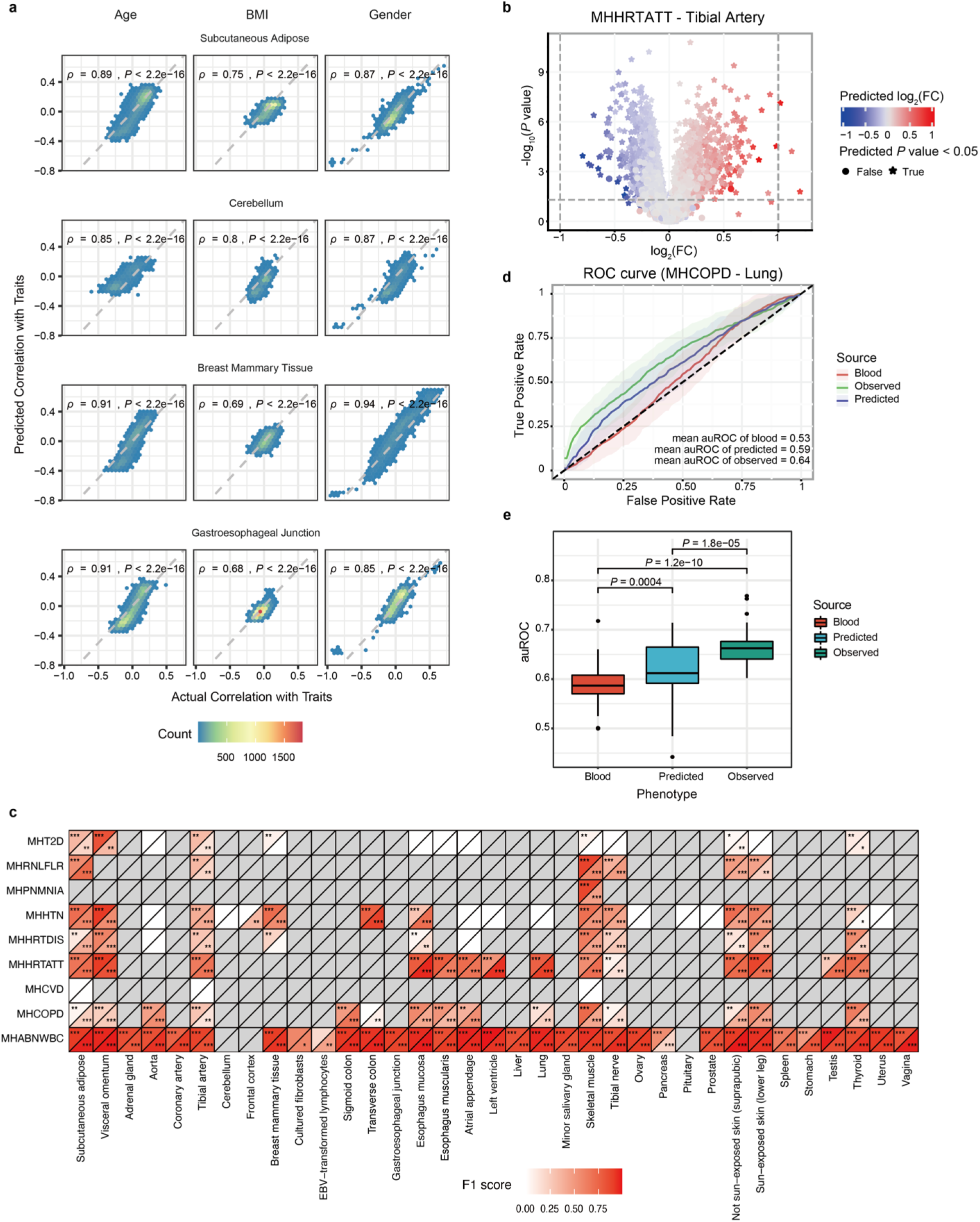
Predicted data preserve personalized biological variations. **a** Hexagonal heatmaps show the relations between the associations of genes with traits (age, gender and BMI) using predicted expression and those using the actual expression within GTEx. Concordance in terms of Spearman’s correlations (Spearman’s *ρ*) between association levels (Pearson’s *ρ*) based on the predicted and actual gene expression are displayed in the plots. Hexagonal cells are coloured according to numbers of genes (count). **b** Comparison of results in differential expression (DE) analysis between the predicted expression and the observed tibial artery expression with heart attack. The position of each point represents the observed results, while the colour indicates the predicted log_2_ fold change. **c** Performance of predicting differentially expressed genes (DEGs) of different diseases in different tissues using predicted data. The number of stars in the grids indicates the overlap significance level of up- or downregulated genes between observed and predicted data (*= significant at 0.05, ** = significant at 1×10^−10^ and *** = significant at 1×10^−100^). The colour represents the level of the F1 score for predicting DE genes. **d** A specific case of receiver operating characteristic (ROC) curves for predicting chronic obstructive pulmonary disease (COPD) status in the observed lung expression, the predicted lung expression, and the input blood expression. **e** Performance of predicting disease status (auROC) of different cases using observed tissue data, predicted tissue data, and input blood data.

Apart from trait-associated changes, we investigated the consistency between tissue-specific disease-related dysregulations in the predicted data and those of the actual data (GTEx). Differential expression analysis (DE analysis) in pGenes was conducted for the combination of each tissue and each disease with at least 50 cases and 50 controls annotated in GTEx, resulting in 117 disease-related tissue-disease pairs across 35 tissues and 9 diseases (with at least 10 differentially expressed genes (DEGs), FDR < 0.05). In these 117 disease-related pairs, 112 pairs exhibited both high sign consistency (sign consistency > 0.7) and high correlation (Pearson’s *ρ* > 0.3) of log_2_ fold changes of pGenes, among which DEGs showed even higher concordance (Supplementary Table 3), suggesting that MTM captured both the direction and the level of the disease-related expression changes. Among 109 pairs with at least 10 observed upregulated DEGs, the predicted DEGs significantly overlapped (hypergeometric test, *P* < 0.05) with the reference DEGs in 93 pairs (F1_median_ = 0.65, F1 score > 0.7 in 44 pairs). For downregulated DEGs, the predicted DEGs significantly overlapped with reference DEGs in 94 pairs (F1_median_ = 0.60, F1 score > 0.7 in 41 pairs) among 112 affected pairs (with at least 10 observed downregulated DEGs, Fig. 4b–c).

To assess whether the predicted tissue expression profiles could be employed in indicating disease status, we next compared the prediction performance of the predicted tissue expression, the actual tissue expression and the input blood expression profiles. Among the 117 disease-related pairs mentioned above, 53 pairs were focused where the actual tissue expression profiles were more informative than those of blood for disease status prediction with *areas under the ROC curve* (auROCs) of at least 0.6. Although less predictive (Wilcoxon paired test, *P* = 1.82×10^−5^) than the actual tissue expression (auROC_mean_ = 0.66), the mean auROCs of the predicted tissue expression of pGenes across the 53 pairs (auROC_mean_ = 0.62) were significantly (Wilcoxon paired test, *P* = 4.03×10^−4^) higher than those of the original blood expression (auROC_mean_ = 0.59), suggesting that predicted tissue expression profiles were informative for disease status (Fig. 4d–e and Supplementary Fig. 8).

### MTM facilitates the identification of tissue-specific disease-associated dysregulations

Since MTM could capture individualized biological variations from any input tissue expression profiles, we further explored the potential application scenarios for MTM. We investigated whether MTM could predict disease-related dysregulations of tissues from blood expression on an external dataset. The external blood expression data were input into MTM to predict the expression profiles of other tissues, which were then used to perform DE analysis and were compared with the GTEx reference (with at least 50 cases and 50 controls). The dataset includes 25 type 2 diabetes (T2D) patients and 33 normal subjects [33]. Among 16 affected tissues (with at least 10 observed up- or downregulated DEGs under FDR < 0.05 within GTEx), the predicted T2D-related expression changes (fold changes) exhibited high concordance with the reference data (Pearson’s *ρ* ranged from −0.49 to 0.76, median = 0.52, Fig. 5a), which were significantly higher (Wilcoxon rank sum test, *P* = 0.03) than those of 11 unaffected tissues (Pearson’s *ρ* ranged from −0.54 to 0.65, median = 0.32).

**Fig. 5.**
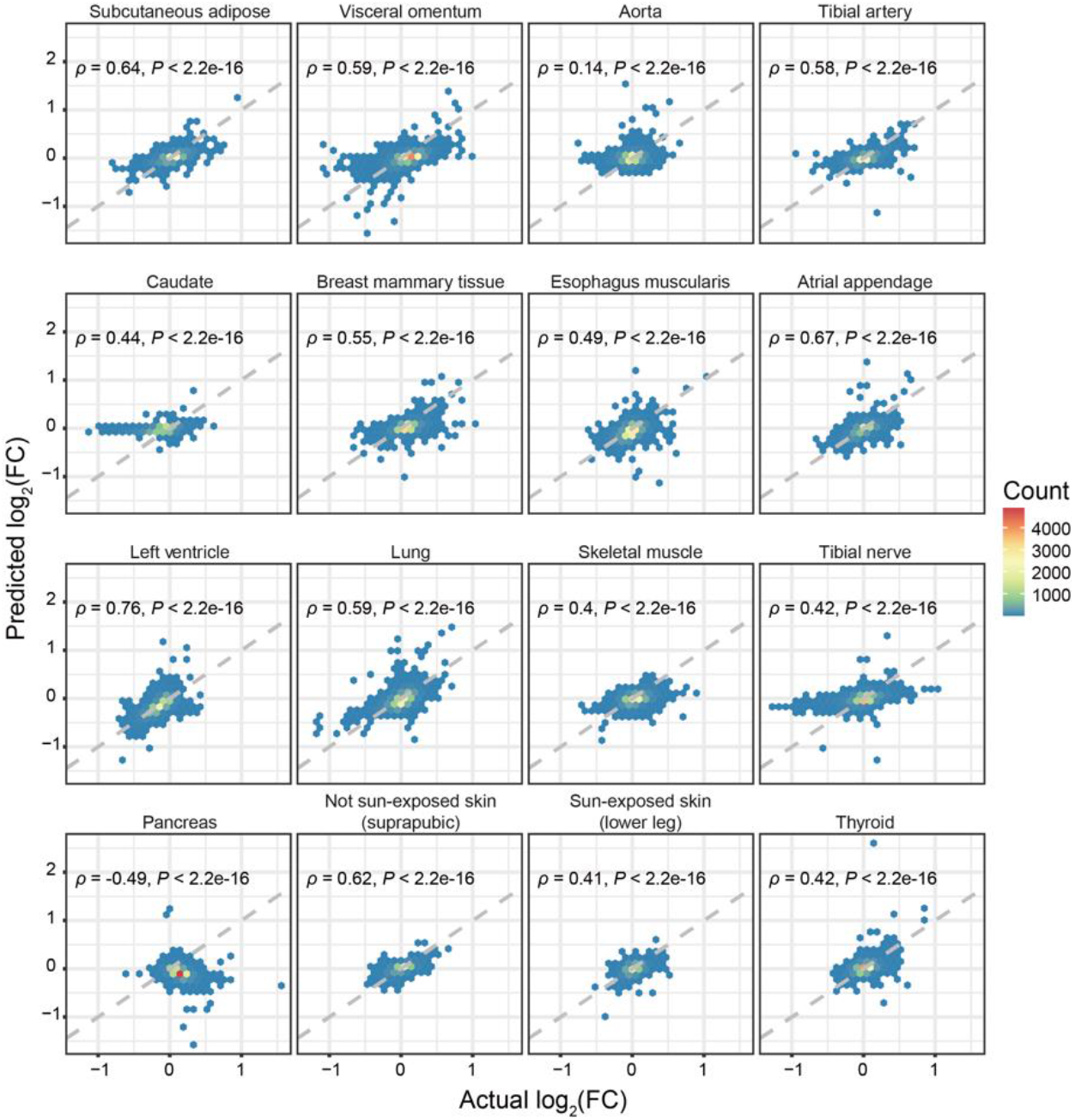
Potential utilities in analysing disease-related gene signatures in external data. Hexagonal heatmaps show the relations between the log_2_ fold changes of the predicted tissue expression (using blood from external T2D patients) and the reference log_2_ fold changes (GTEx T2D subsets). Hexagonal cells are coloured according to numbers of genes (count).

## Discussion

Predicting tissue transcriptome profiles from biological information of peripheral ‘surrogate’ samples has emerged as an effective alternative with potential utility in areas of biomedical science, especially for circumstances when tissue samples are unavailable. In this work, we have proposed MTM, a deep-learning-based multi-task learning framework that enables the prediction of individualized tissue gene expression profiles from any available gene expression of the same individual using a single unified model. Through the nonlinear representation capability of neural networks and by fully leveraging information from all tissues of the same individual via the multi-task learning architecture, MTM not only bypasses the need to establish an independent model for one mapping direction (such as blood to one brain area), but also substantially enhances the performance on both the overall sample levels and fine-grained gene levels. In particular, MTM identifies more predictable genes with well-preserved personalized physiological and pathological characteristics, providing a novel and valuable tool for tissue transcriptome-based biomedical investigations from a more comprehensive and systemic perspective.

The substantial improvement in performance for MTM may be attributed to several factors. First, it is commonly recognized that an individual’s gene expression levels in different tissues are inherently linked by shared biological foundations, which could be decomposed into the static part that shares the same genetic material across tissues (i.e., GReX) and the dynamic part that is related to the temporal and spatial factors of the individual, including physiological states as well as environmental and other factors. The multi-task learning design of MTM encourages the exploitation of both the static and dynamic intrinsic relevance in all tissues from the same individual, enabling the model to extract and refine decoding rules with biological patterns and to produce better individualized representations instead of solely twisting data, thus leading to improved predictive performance, especially in tissues with smaller sample sizes. Interestingly, MTM performed worst in the two cell lines, with the lowest numbers of pGenes being captured, indicating that the integrity of internal information for the individual was lost in the cell lines that were cultured or transformed *in vitro*. Second, MTM may be more capable of capturing complex gene–gene interactions with nonlinearity and multi-task components. Last but not least, predictive information, including subtle signals, is preserved to the greatest extent in MTM, since dimension reduction is not required for transcriptomic data in large-scale neural networks driven by the backpropagation algorithm [34]. With these properties, the framework of MTM may serve as an effective paradigm in similar biomedical scenarios where complex intrinsic relationships underlie multiple entities, such as other large-scale omics data of different tissues or cells from the same individuals.

The high accuracy of MTM for cross-tissue expression prediction with well-preserved individualized biological variations supports its potential for biomedical applications, especially in tissue-specific biomarker discovery in liquid biopsy. However, there are still gaps that remain to be filled. A major challenge is to remove the heterogeneity in real data to align with the fixed develop sets for deployed machine learning models in practice, which imposes difficulties on existing data integration methods that cannot keep the develop sets intact. Another challenge is that the multi-task learning architecture of MTM highly depends on large-scale computational resources, which limits the incorporation of additional omics data. With the future advances of computational resources and data integration algorithms, as well as new incoming data to support continuous learning, MTM will promote real-world translational applications, such as decoding molecular mechanisms and mining clinical biomarkers.

## Conclusions

In this study, we propose a deep-learning-based multi-task learning framework, MTM, to predict individualized tissue gene expression profiles from any available gene expression of the same individual using a single unified model. Comparisons to the two representative approaches, MTM show the superior performance at both the sample level and the gene level, with a larger proportion of predictable genes. Ablative study confirms that jointly leveraging cross-tissue information provides improvements beyond the modeling power of nonlinear neural networks, which might be achieved by exploiting individualized representation through our multi-task learning framework. Phenotype association analysis suggests that the predicted expression profiles preserve the individualized biological variations well, including trait-related variations and disease-related dysregulations. In summary, our work proves multi-task learning to be an effective strategy to utilize tissue-shared intrinsic biological relevance in the prediction of cross-tissue gene expression profiles.

## Methods

### Data preparation for MTM

The gene expression data from human tissues were downloaded from the v8 release of the Genotype-Tissue Expression (GTEx) portal (https://www.gtexportal.org/) [32]. The metadata, including the subject-level and sample-level information, were obtained through the database of Genotypes and Phenotypes (dbGaP) (https://www.ncbi.nlm.nih.gov/gap/). Individuals whose blood RNA-seq and genotype data were available were included in our study. The expression values (transcripts per million, TPM) of 19,291 protein-coding genes from 17,329 samples (49 tissues with sample sizes greater than 50) across 948 individuals were used for model development. The tissue expression profiles were randomly split into the training set (80%) and the validation set (20%) based on individual labels. Z-score standardization was conducted within each tissue for the training data, and then the recorded scaling factors were used for the validation data to prevent information leakage.

### Description of MTM

#### Key elements

The MTM architecture consists of an encoder, a generator, a discriminator, and a mapping network. The encoder (*E*) embeds expression profiles into a shared latent space, resulting in personalized representations, i.e., latent codes. The generator (*G*) is symmetrical to the structure of the encoder, through which the latent codes were mapped into tissue-specific expression spaces in the form of individualized tissue expression profiles. The discriminator (*D*) is employed to identify the mismatches of the predicted distributions and the actual distributions to send feedback signals for the generator to improve the prediction in an adversarial manner [35]. The auxiliary mapping network (*M*) assists *E* in learning a smooth space [36]. All *E, G* and *D* networks are multi-task networks equipped with TCMs, which were designed to route and transform the data through tissue-specific or shared connections. The TCM consists of three parts: a shared fully connected layer and a learnable instance-level affine transformation conditioned on tissue labels followed by a leaky rectified linear unit (Leaky ReLU) to introduce nonlinearity. The detailed structure of the MTM is described in Supplementary Fig. 9–13.

#### Objectives

The generator (*G*) translates latent codes from a shared latent space (*c* ∈ 𝒲) to predict the expression profiles conditioned on the target tissue *t, G*(*c*|*t*) → *x*_*t*_. The latent codes could either be reference-guided by *E* (*E*(*x*|*s*) → *c*) or transformed from random Gaussian noise 𝒵 ∼ 𝒩(0,1) by 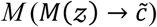. To enforce *G* to utilize the latent code 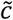 when predicting the expression profile 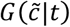, we employed the L1 reconstruction loss:

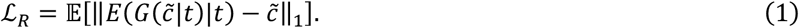

To minimize the divergence between the distributions of the actual expression profiles and of the predicted ones (*G*(·)), a training strategy from a generative adversarial net (GAN) is used, where *D* tries to distinguish the generated samples from real samples, while *G* tries to fool *D*. Instead of the original GAN, which minimizes the Jensen–Shannon divergence [35], we used an adversarial objective function in the form of the hinge loss for robust training [37, 38]:

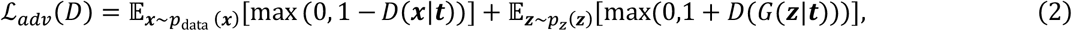

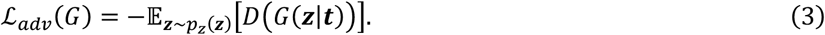

To achieve individualized tissue expression prediction, given the expression profile of the source tissue *x*_*s*_ from a specific individual, we use the L1 loss to penalize the difference between the predicted expression profiles *G*(*E*(*x*_*s*_|*s*)|*t*) and the actual expression profile of the target tissue *x*_*t*_:

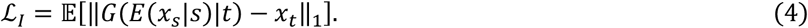

To ensure that the predicted expression profile properly preserves the individual invariant characteristics, a cycle consistency loss is employed:

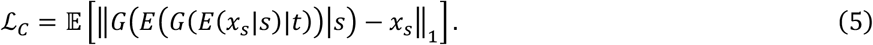

The final objective function is as follows:

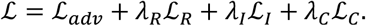

#### Training hyperparameters

The data were split into mini-batches of 256 samples for training. The Adam optimizer [39] was employed to optimize the network parameters using the following hyperparameters: learning rate = 0.0005, β1 = 0.5 and β2 = 0.9. The number of maximum training epochs was set to 200. All models were trained using the PyTorch framework (version 1.10.2) [40] on Tesla V100S graphics processing units (NVIDIA).

### Description of intermediate model S3 for ablative experiments

The S3 model consisted of a set of nonlinear neural networks for each tissue based on simple multilayer perceptrons. For each tissue, a fully connected neural network with Leaky ReLU activations was built to predict the expression profiles of the target tissue from the expression of a specific source (blood). The number of maximum training epochs was set to 1000. Other training hyperparameters were kept the same as those of MTM.

### Performance evaluation

The prediction quality was indicated by the Pearson correlation coefficient (Pearson’s *ρ*) between the predicted and observed expression data. The sample-wise accuracies across all genes were calculated on Z-score standardized data to highlight the individual variance in each tissue. The gene-wise accuracies refer to Pearson’s *ρ* of each gene across individuals for each tissue. Predictable genes were defined as genes with gene-wise *ρ* > 0.3 between predicted and observed expression in each tissue. Accuracy indicators in this work were reported in a fivefold cross-validation (CV) manner. For fair comparison, the performance of S1, S2 and S3 was evaluated in a CV manner using the same inputs as were used with MTM (blood expression data from the same training and validation samples).

### Exploration of the characteristics of predictable genes

#### Tissue similarity

Similarities between tissues were measured with Pearson’s *ρ* between the mean expression levels across genes (log2-scaled) of each tissue. When comparing certain pairwise statistics (x1) of tissues with the tissue similarities (x2), element-wise values were flattened to two arrays (X1 and X2), and then Spearman’s correlation coefficient between the two was calculated.

#### Expression levels

In each tissue, the Wilcoxon rank sum test was used to compare the mean expression levels of the top 25 percentiles of predictable genes (highly predictable genes) with those of the control groups, with the control groups defined as the bottom 25 percentiles of predictable genes (unpredictable genes).

#### Conservation

The conservation distributions of genes in terms of PhastCons scores were extracted with DeepTools [41] from the 100-way vertebrate species alignment downloaded from the UCSC (University of California Santa Cruz) genome browser [42, 43]. The conservation levels of the highly predictable genes were compared with those of the unpredictable genes in each tissue (Wilcoxon rank sum test).

#### Connectivity

The degree of connectivity distribution for each gene was extracted from the downloaded PIN. Then, the average degrees of the two groups of genes (highly predictable genes and unpredictable genes) were compared. The average degrees of the highly predictable genes were compared with those of the unpredictable genes in each tissue (Wilcoxon rank sum test).

### Exploration of the characteristics of the intermediate features of MTM

Personalized representations from the latent space of the MTM encoder were extracted across different individuals and tissues and then used to calculate pairwise similarities (Pearson’s correlation coefficients). Intraindividual codes refer to latent codes of different tissues from the same individual, while interindividual codes refer to latent codes from different individuals.

Decoding paths of the data flow from latent codes to tissue expression spaces refers to activations of different layers through the MTM generator. The decoding paths of each layer from an individual’s code from blood to different tissues were used to calculate pairwise similarities (Pearson’s correlation coefficients). Spearman’s correlation coefficients between the pairwise similarities of the decoding paths and the pairwise tissue similarities were calculated to determine the extent to which the decoding rules reflected biological patterns.

### Phenotype-related analysis

For traits including age, gender and BMI, Pearson’s correlation coefficients were used to measure the associations of each gene with traits in each tissue. Then, Spearman’s correlation coefficients between the predicted and actual (GTEx) associations with traits were calculated to determine the extent to which the predicted tissue expression preserved the trait-related expression changes.

For disease-associated dysregulations, the sign consistency and Pearson’s correlation coefficient between log_2_ fold changes across genes in predicted and actual data were used to measure the overall concordance of the prediction. Differential expression analysis was conducted using the nonparametric Wilcoxon rank sum test. When comparison of differentially expressed genes (DEGs, under FDR < 0.05) between predicted and actual data was applicable, hypergeometric tests were conducted to examine the overlap significance, and the F1 scores, which convey the balance between precision and recall, were used to measure the performance of identifying DEGs.

For disease status prediction, we built least absolute shrinkage and selection operator (LASSO) models with scikit-learn (version 0.24.2) [44] for each disease-tissue pair in the predicted expression, the actual expression and the input blood expression. The values of the *area under the ROC curve* (auROC) were used to indicate performance in a fivefold CV manner with 50 independent repetitions.

### Exploration of MTM applications on external datasets

The raw sequencing data (fastq files) of blood expression profiles from GSE184050 [33] were downloaded, and the gene expression values of the 19,291 protein-coding genes were quantified according to the pipeline provided by the GTEx Consortium [32]. To reduce batch effects, we simply aligned the mean and standard deviation of each gene in each tissue to those of the GTEx data. For external dataset, the MTM was trained in a hold-out manner, with 80% of the data used for training and the remaining 20% for testing.

The standardized blood expression profiles of GSE184050 (type 2 diabetes, T2D) were input into MTM to predict the tissue expression profiles of corresponding individuals. Then, Pearson’s correlation coefficients between log_2_ fold changes across genes in predicted data and reference data (case and control subjects for T2D in GTEx) were used to measure the overall concordance of the predicted disease-related dysregulations.

